# A human mesenchymal spheroid prototype to replace moderate severity animal procedures in leukaemia drug testing

**DOI:** 10.1101/2022.07.07.499143

**Authors:** Aaron Wilson, Sean Hockney, Jess Parker, Helen Blair, Deepali Pal

## Abstract

Patient derived xenograft (PDX) models are regarded as gold standard preclinical models in leukaemia research, especially in testing new drug combinations where typically 45-50 animals are used per assay. 9000 animal experiments are performed annually in leukaemia research with these expensive procedures being described as moderate severity, meaning they cause significant pain, suffering and visible distress to animal’s state. Furthermore, not all clinical leukaemia samples engraft and when they do data turnaround time can be between 6-12 months. Heavy dependence on animal models is because clinical leukaemia samples do not proliferate in vitro. Alternative cell line models though popular for drug testing are not biomimetic – they are not dependent on the microenvironment for survival, growth and treatment response and being derived from relapse samples they do not capture the molecular complexity observed at disease presentation. Here we have developed an in vitro platform to rapidly establish co-cultures of patient-derived leukaemia cells with 3D bone marrow mesenchyme spheroids, BM-MSC-spheroids. We optimise protocols for developing MSC-spheroid leukaemia co-culture using clinical samples and deliver drug response data within a week. Using three patient samples representing distinct cytogenetics we show that patient-derived-leukaemia cells show enhanced proliferation when co-cultured with MSC-spheroids. In addition, MSC-spheroids provided improved protection against treatment. This makes our spheroids suitable to model treatment resistance – a major hurdle in current day cancer management

Given this 3Rs approach is 12 months faster (in delivering clinical data), is a human cell-based biomimetic model and 45-50 animals/drug-response assay cheaper the anticipated target end-users would include academia and pharmaceutical industry. This animal replacement prototype would facilitate clinically translatable research to be performed with greater ethical, social and financial sustainability.

**Research Highlights:** *Scientific Benefit:* A 3D spheroid-based approach for ex vivo co-culture of clinical leukaemia samples for further investigation into cancer biology such as blast-niche interactions, blast proliferation and treatment resistance

*3Rs Benefit:* To replace moderate severity animal procedures in leukaemia research and drug testing

*Practical Benefit:* 3Rs approach that yields drug response data quickly and is more ethically, socially and financially sustainable than its in vivo counterparts

*Current applications:* Exploration of leukaemia biology such as blasts proliferation, blast-niche interactions, niche-impacted treatment resistance and obtain drug response data

*Potential applications:* Extend the approach to include other haematological cancers as well as bone cancers.

## Introduction

Leukaemia management is hindered by two chief obstacles: 1. Treatment still involves toxic chemotherapy drugs(*1, 2*) 2. 15-20% of patients go into relapse at which point the disease can be treatment resistant(*3*). The nature of relapse disease highlights the need for safer and improved treatments such as targeted therapy. While many novel therapeutics have been identified, high drug attrition rates have been a major hindrance against anti-cancer drug development. 95% of bench side drugs that make it to phase 1 clinical trial never reach patients(*4-6*). Three key areas(*5*) have been identified that must be addressed in order to reduce attrition rates: the first being an increase in scientific collaboration and the second being that trials must be tumour biology driven for the drug to be more therapeutically effective. The third issue highlights the lack of robust and translatable preclinical cancer models that could be used to screen out many unsuccessful therapeutics before going to trial(*5*).

The gold standard preclinical model in leukaemia research is a patient derived xenograft model(*7-9*). Recent Home Office annual returns show that 195,407 mice are used annually in cancer research. Indeed, cancer research remains the most common applied sciences area to use animal procedures. Given leukaemia research is heavily dependent on mouse models combining these metrics with financial data from major cancer funders in the UK suggest that 5% of these mice or at least 9000 mice are used in leukaemia research every year. Although leukaemia is a very aggressive disease, leukaemia cells do not maintain viability outside the body(*9*). This has led to the development of patient-derived xenograft (PDX) models where leukaemia cells are engrafted into the BM of immunocompromised mice. PDX models are currently the most clinically relevant models in blood cancer preclinical studies. Animal(mouse) experiments of moderate severity are used to amplify patient-derived leukaemia cells, to study leukaemia biology and perform translational studies(*8, 10*). Patient-derived leukaemia cells can take anywhere between 2-12 months to engraft in mice before any preclinical drug testing can be carried out, hence these mice experiments are very time consuming. In addition, most of these procedures are considered to be moderate severity and often involve mice developing significant adverse effects such as weight loss, fur ruffling, back hunching as well as increased white cell counts.

The pipeline for preclinical testing as documented in literature is: 1. Toxicity analysis to determine no observed adverse effect limit (NOAEL) / maximum tolerated dose (MTD): 15-20 mice per drug(*11*). 2. Pharmacokinetic studies to determine drug maximum plasma/tissue levels and kinetics of drug clearance:15-30 mice per drug(*12*). 3. Drug efficacy studies and pharmacodynamics assays on tumour cell response: 5-10 mice needed per drug treatment group(*13*). Per drug group a total of 40-50 animals are needed for single drug assays and twice as many for drug combination assays(*11-13*). In addition, in vitro cancer models preceding in vivo validation experiments rely on cell lines that show limited in vivo predictability. Cell lines retain cytogenic characteristics of ALL but being derived from relapse do not represent the molecular complexity of disease at presentation. Cell lines have also artificially adapted to grow in suspension culture which eliminates a key feature of leukaemia biology: microenvironment and its role in treatment resistance. Furthermore these models are restricted by limited in vivo predictability and often lead to in vitro to in vivo drug attrition rates. This in turn results in unnecessary animal experiments which could have been avoided for these failed drug candidates, i.e., false positive data from cell line models.

In a preceding pilot study, we developed an ex vivo 2D co-culture platform, the purpose of which has been to conduct preclinical drug screening using patient-derived leukaemia cells thereby reducing dependence on cell lines and consequently minimising in vitro to in vivo drug attrition rates.

ALL is a disease primarily of the bone marrow (BM) microenvironment (or leukemic niche), a compartment within the bone composed of cell types including mesenchymal stem cells (MSC), osteoblasts, osteoclasts, endothelia, immune cells and supporting perivascular cells. Difficulty in growing and maintaining patient derived samples once they removed from the patient alludes to the vital role that the niche plays in cancer biology. Study into blast-niche interaction has unveiled some of the supportive roles the niche plays in disease, including cell-cell signalling, growth factors, cell adhesion and evidence has shown the niche providing an element of chemoprotection(*14, 15*). The niche particularly mesenchyme-blast interaction gives rise to two lymphoblast subpopulations, the first being a slow cycling, dormant population and the second being actively cycling disease propagating blasts(*16*). Many treatments are effective in reducing the populations of active cycling blasts; however, it is reported that the dormant populations can survive treatment, seeking refuge within the niche. These dormant cells can later shift into the cycling state post treatment, leading to relapse disease(*16, 17*).

Targeting of niche-blast interactions opens a door into a variety of potential therapeutic approaches. An example would be CXCL12, a chemokine secreted by stroma cells and upregulated in disease, it was discovered that CXCL12 induces a non-cycling state within LSCs, reducing the effectiveness of tyrosine kinase inhibitors. Mesenchymal cell CXCL12 knockout reversed this change and caused a phenotypic switch into disease propagating cycling blasts, temporarily increasing disease load but allowing once chemo resistant blasts to be treated(*18*). Here we model the leukaemia stromal niche using human BM-MSC-spheroids.

Recent advances in medicine have utilised 3D models as miniature in vitro representations of specific organs, they are produced in 3D and retain a realistic microanatomy, allowing a representational human model(*19*). 3D models recapitulate their respective organ function and have been proven to be useful models of human disease, particularly in cancer research(*20*). 2D models cannot recapitulate the leukemic niche or the complex and varied interactions that influence leukemic cell growth(*21*). Current medicine is making use of organ-on-a-chip systems which have led to advances in liver and lung drug screening, providing more accurate toxicological reports than 2D systems(*19*). The complex and dynamic nature of ALL and the bone marrow mesenchymal microenvironment makes it an ideal candidate for a more representative drug screening platform, this paper aims to do this by developing mesenchymal spheroids, which will support the culture of patient derived leukaemia samples allowing patient-specific therapeutic investigation(*22*).

## Materials and methods

### Materials

| General |  |
| --- | --- |
| Material | Manufacturer |
| Trypsin-EDTA Solution 10X | Sigma-Aldrich, Dorset, UK |
| Phosphate-Buffered Saline | Thermo Fisher Scientific, Hertfordshire, UK |
| Gentle Cell Dissociation Reagent | Stem Cell Technologies, Stem Cell, UK |
| Heat Inactivated Foetal Bovine Serum | Thermo Fisher Scientific, Hertfordshire, UK |
| Matrigel HESC-Qualified Matrix | Corning, Kennebunk, USA |
| Low-glucose Dulbecco's Modified Eagle's Medium | Sigma-Aldrich, Dorset, UK |
| Mesenchymal Stem Cell Culture |  |
| Material | Manufacturer |
| Low-glucose Dulbecco's Modified Eagle's Medium | Sigma-Aldrich, Dorset, UK |
| Heat Inactivated Foetal Bovine Serum | Thermo Fisher Scientific, Hertfordshire, UK |
| Penicillin/Streptomycin | Sigma-Aldrich, Dorset, UK |
| Basic FGF | Sigma-Aldrich, Dorset, UK |
| L-Glutamine | Sigma-Aldrich, Dorset, UK |
| Matrigel HESC-Qualified Matrix | Corning, Kennebunk, USA |
| Leukaemia Blast Co-Culture |  |
| Material | Manufacturer |
| SFEM, Serum-Free Medium for Culture and Expansion | Stem Cell Technologies, Stem Cell, UK |
| Organoid formation |  |
| Material | Manufacturer |
| AggreWellM 400 24-well Plate | Stem Cell Technologies, Cambridge, UK |
| AggreWell Rinsing Solution | Stem Cell Technologies, Cambridge, UK |
| Costar Ultra-low Attachment 6 Well Plate | Corning, Kennebunk, USA |
| Real-Time PCR Analysis |  |
| Material | Manufacturer |
| QIASHredder | Qiagen, Manchester, UK |
| Beta-mercaptoethanol | Sigma-Aldrich, Dorset, UK |
| RNEasy Micro Kit | Qiagen, Manchester, UK |
| RNase-Free DNase Set | Qiagen, Manchester, UK |
| RT2 First Strand Kit | Qiagen, Manchester, UK |
| RT2 Profiler PCR Array | Qiagen, Manchester, UK |

Reagents table.

### Methods

#### Ethical approval

Patient-derived leukaemia blasts were obtained from the Newcastle Biobank (REC reference number 07/H0906/109+5). All samples were obtained following written informed consent. All animal studies were carried out in accordance with UK Animals (Scientific Procedures) Act, 1986 under project licence P74687DB5 following approval of Newcastle University animal ethical review body (AWERB).

#### 2D Niche-Blast co-culture

Patient derived ALL samples were cultured on 2D monolayers of MSC cells. MSC were seeded on Matrigel coated 48 well plates at 2×104 cells/0.5mL/2cm2 in their respective media. After 24 hours, the media was removed and the cells rinsed with 1mL SFEMII, Leukaemia blasts were then plated at a density of 0.25×105 cells/mL per well suspended in SFEMII.

#### 3D culture of BM-MSC

MSC cells were transferred to a 6-well low adhesion plate for suspension culture and cultured in 2 mL MSC media 3D for 48 hours, after which, MSc spheres were seen to form. Spheroids obtained thus were irregular in shape therefore Aggrewell400 plates were used to create uniform MSC spheres. Aggrewell plates were prepared by rinsing each well with 0.5mL of Aggrewell rinsing solution, spinning at 1500g for 5 minutes and aspirating. MSC cells were seeded at varying concentrations depending on desired sphere size. Cells were split using 1x trypsin solution and seeded in single cells on Aggrewell plates at desired concentration in MSC media. Aggrewell plates were spun at 100g for 3 minutes to ensure all cells reached the bottom of the microwells. After 48 hours, MSC spheroids were ready for harvest and transferred for culture onto low adhesion plates

#### 3D BM-Blast co-culture

Blasts were co-cultured with the MSC spheres in SFEMII at 2×105 cells/mL in 5ml of media in low adhesion 48 well plates.

#### Dexamethasone 3D drug assay

Investigation into DEX mediated treatment resistance was carried out through cell fate tracing. 10 million blasts were span at 500g and resuspended in 10 mL of 5uM CellTrace violet, incubated for 20 minutes at 37 degrees Celsius, 1 mL FBS was added and cells were span once more at 500g for 5 minutes and resuspended in fresh SFEMII. Cells were then plated on 3D MSC spheres in 48 well plates as described above. 5nM Dexamethasone was added to desired wells and blasts were cultured for 5 days.

#### Results

#### Development and validation of BM-MSC spheres

We first show that BM-MSC/leukaemia 2D co-culture platform is able to generate drug dose response data that is consistent with existing in vivo data (Fig 1.). We further perform pilot studies where we show that such ex vivo co-culture platforms have ability to bring about significant replacement in animal procedures (Fig 2). In order to fully recapitulate the complexity of the leukaemia niche, a 3D Model with improved biomimicry is required. 3D BM-MSC-spheroids were created by subculturing cells onto low adhesion non-tissue culture treated plates (Fig 3 a-b). These spheroids were tested for expression levels of their respective niche markers using QT-PCR. It was found that both niche sphere types expressed mesenchymal markers CD90, CDH2 and Nestin (Fig. 3 c) similar to their 2D counterparts.

**Figure 1.**
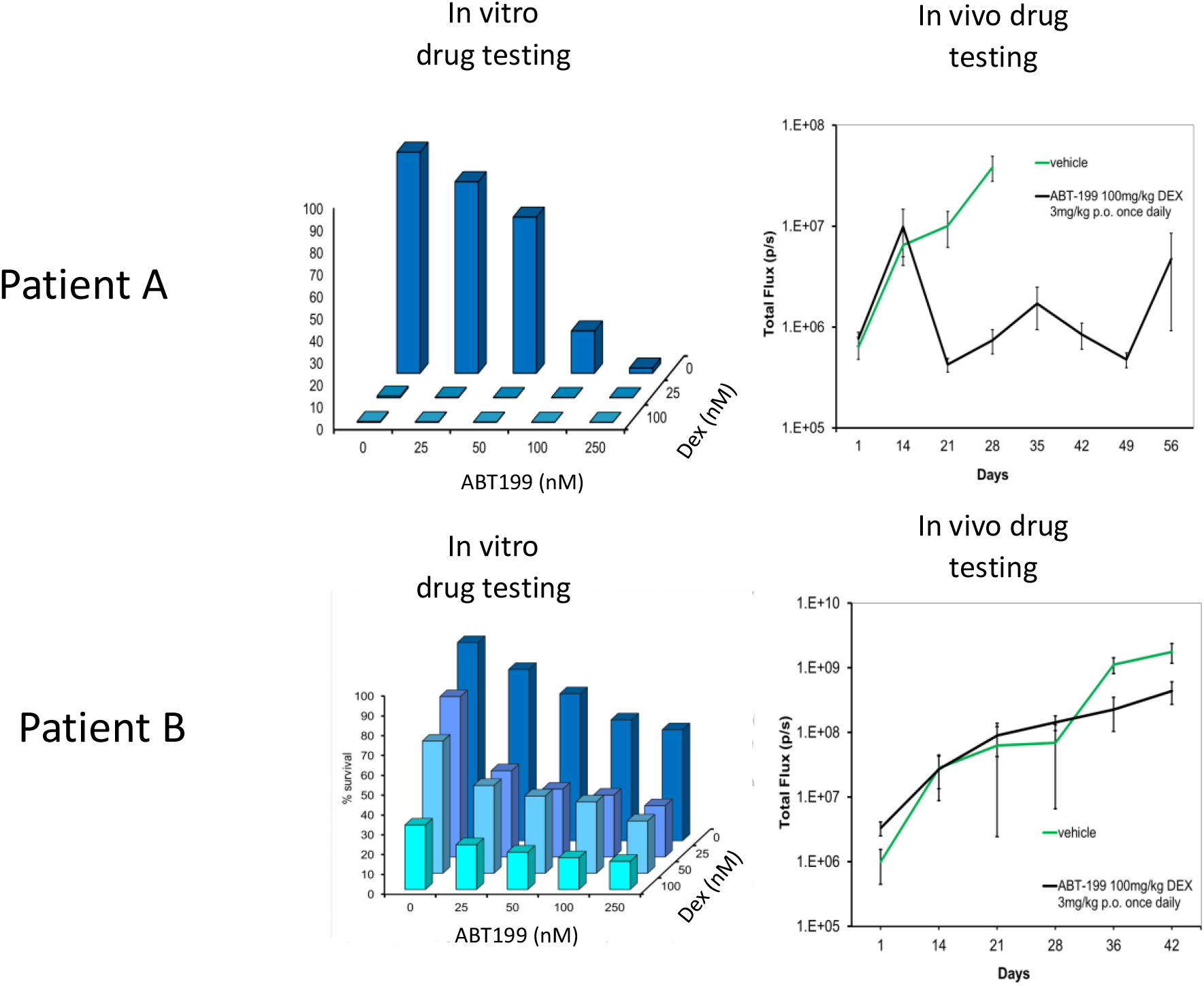
2D *in vitro* platfrom delivers drug response data with high *in vivo* predictability to minimise in vitro to in vivo drug attrition rates. Leukaemia cells from patient A is a responder to ABT199 and Dexamethasone drug combination in vitro and in vivo. On the other hand leukaemia cells from patient B do not respond to this drug combination in vitro and subsequently poor response is also noted in vivo.

**Figure 2.**
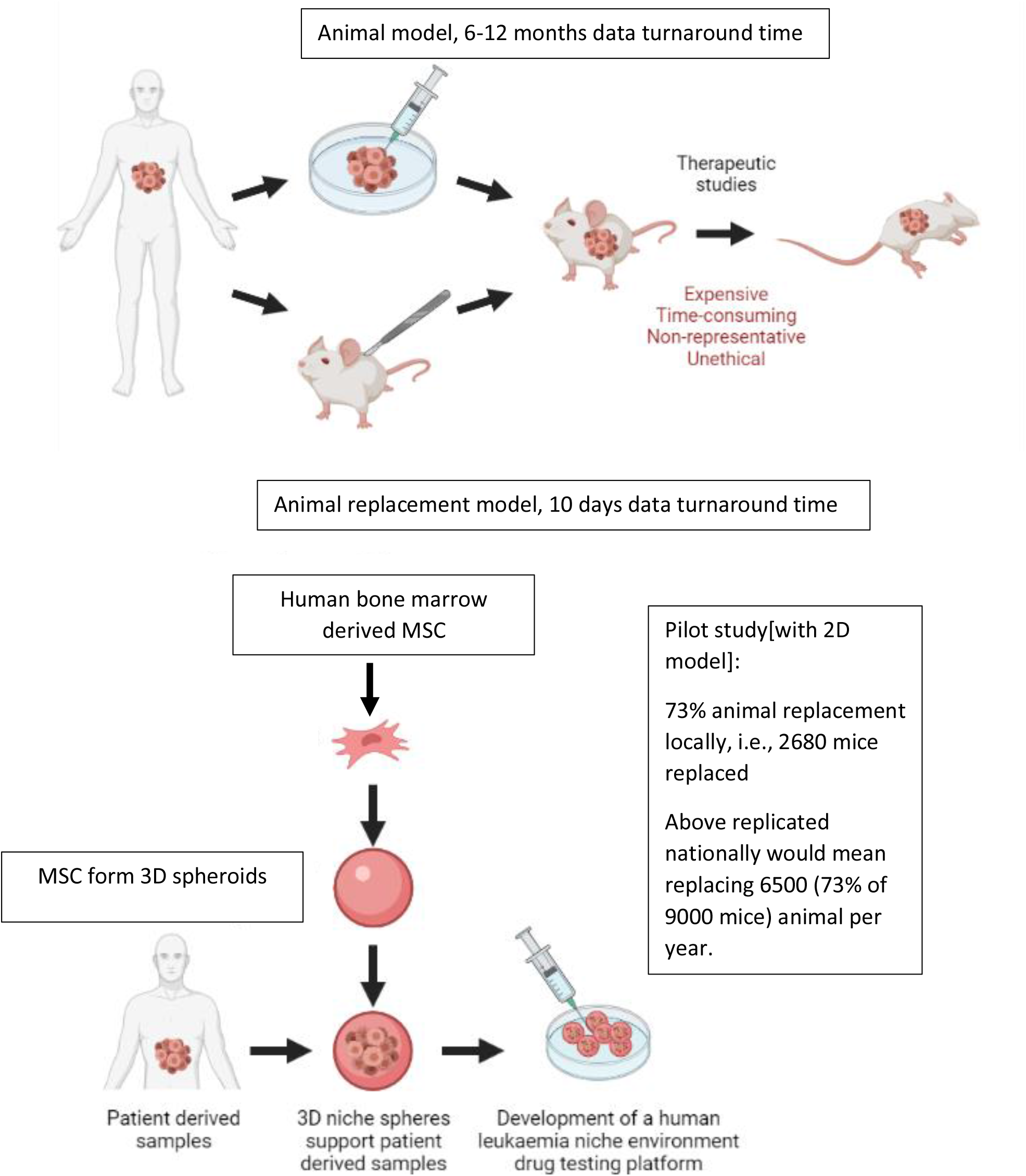
Comparison of existing in vivo preclinical leukaemia models with newly developed ex vivo animal replacement organoid models. Key benefits of the animal replacement organoid models include species specificity and consequently higher biomimicry.

**Figure 3.**
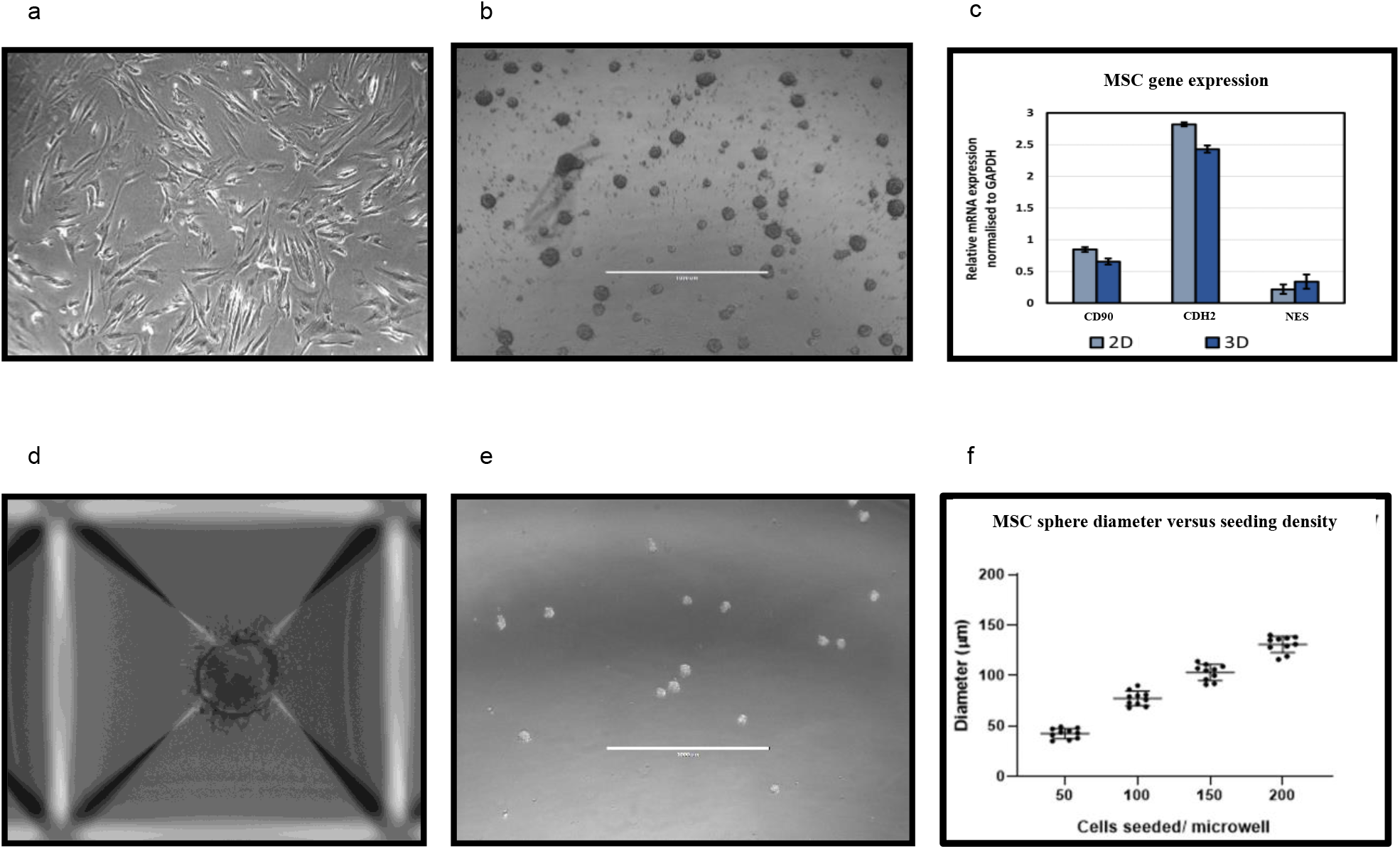
Advancing the 2D ex vivo platform to develop 3D spheroid models. Development and validation of MSC spheroid cultures through 3D suspension culture. Aggrewell plates allow for the creation of uniform 3D spheroids from single cell suspension of the desired cell-type. SD from three independent experiments are plotted as error bars. *2D (a) and 3D MSC (b) after 96 hours of growth. (c) mesenchymal marker levels quantified through qPCR and normalised to GAPDH*. **(d**,**e)** *MSC spheroids obtained through Aggrewell plate culture. (f)Cell seeding densities directly correlate to size of spheroids obtained. Data captured following 48 hours of seeding*.

Aggrewell400 plates are designed for higher throughput and more uniform production of MSC spheres. Each well on these plates house 400 conical microwells, funnelling single cells and encouraging them to conglomerate into spheroids (Figure 3 d). These microwells were utilised to create uniform niche spheres allowing standardisation of the technique when producing assays involving 3D spheres (Figure 3 e). A direct correlation was observed with seeding cell density and size of spheroids with higher seeding densities resulting in generation of more uniformly sized spheroids (Fig.3f).

Patient-derived leukaemia cells co-cultured with MSC show superior proliferation and enhanced treatment resistance

ALL blasts from three different clinical samples were co-cultured with BM-MSC in routine 2D cultures and with 3D spheroids. Cell proliferation was monitored over a 5 day period through tryphan blue cell counts. We observed that when co-cultured with BM-MSC-spheroids the blasts showed 2-fold higher cell proliferation compared to 2D co-cultures (Fig. 4.a-c). Next, we repeated the co-cultures under treatment with the steroid Dexamethasone which is used to treat ALL. We observed that blasts showed significant reduction in sensitivity to Dexamethasone in 3D MSC organoid co-cultures compared to 2D co-cultures (Fig.4.d-f). These data show the advantage of 3D organoid based co-culture systems over routine 2D cultures in two respects: 1. Achieving blast cycling which is essential for the action or testing of majority of anti-cancer treatments 2. Modelling treatment resistance in the laboratory – a major current day clinical challenge in cancer management. Finally, we perform a costs analysis and show improved financial sustainability in our ex vivo organoid platform compared to animal models(Fig 5).

**Figure 4.**
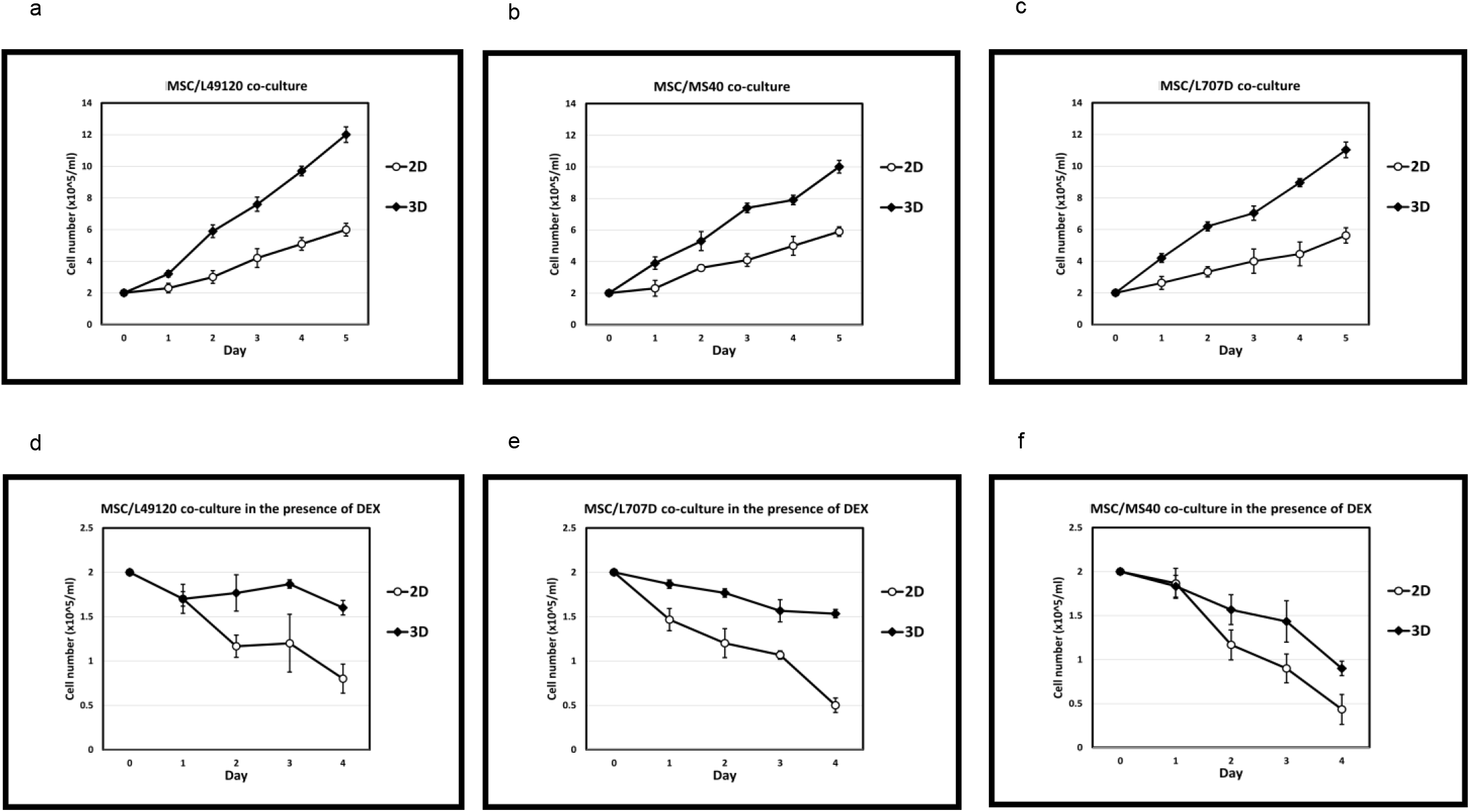
Co-culture of patient derived leukaemia samples on both 2D and 3D MSC +/-treatment pressure. N= 3 Error bars = SD. *Increased growth kinetics of L49120 (a,d), MS40 (b,e) and L707D (c,f) across a 5 day period on 3D MSC feeder cells compared to a 2D monoculture. Dexamethasone treatment is less effective in a 3D microenvironment (d,e,f)*

**Figure 5.**
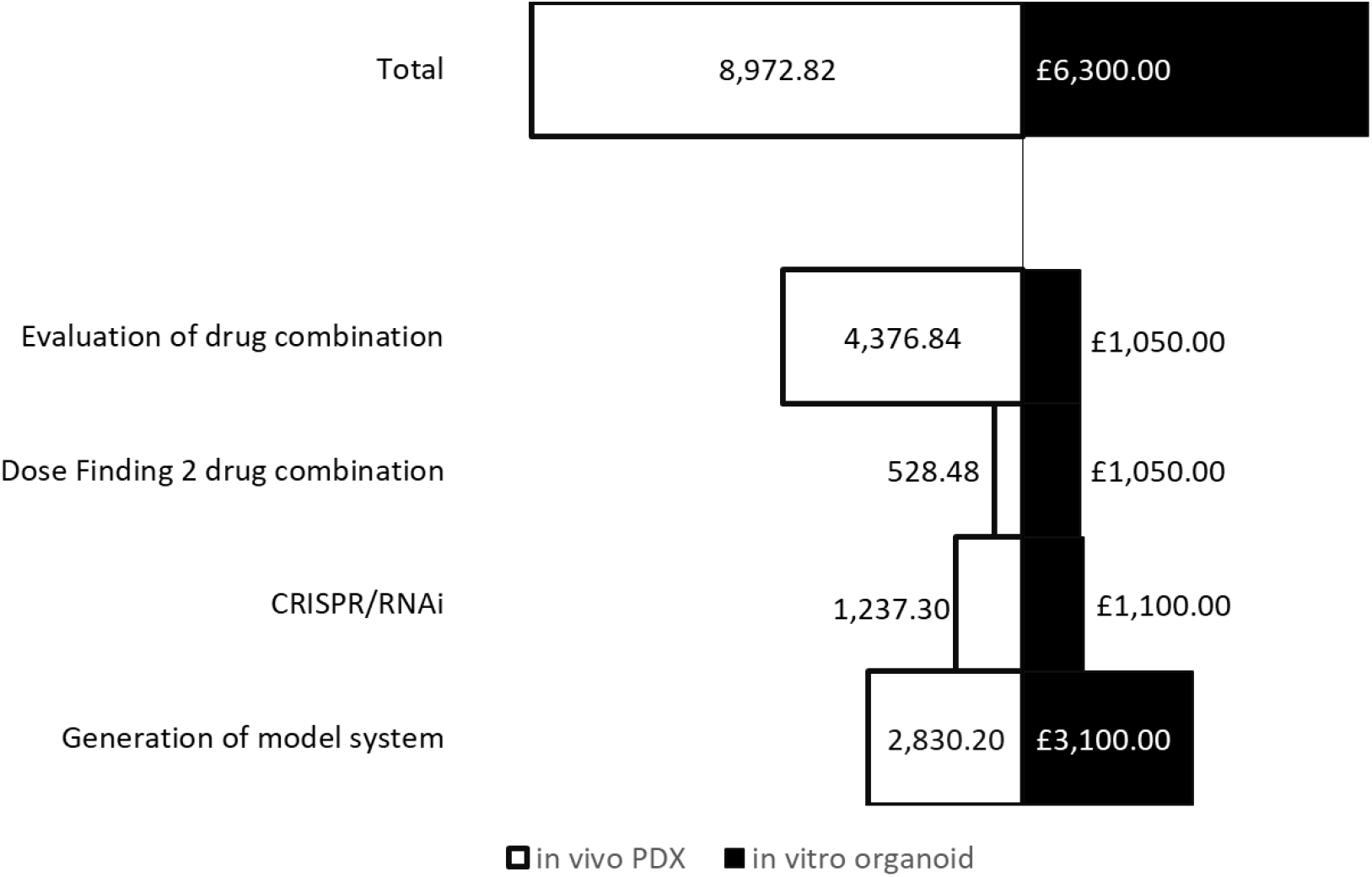
A comparative costs analysis between 3D ex vivo and in vivo preclinical models show an approximate 30% costs benefit and consequently increased financial sustainability when using ex vivo organoid models.

## Discussion

Relapse is still a major concern within the treatment of leukaemia, relapse disease is often chemo resistant, meaning many therapies cease working in patients with relapse disease leading to an increased mortality(*3*). Differing cytogenic abnormalities in this disease are known and patient stratification and precision medicine are implemented in an effort to treat individual disease in the most effective way possible, allowing weaknesses in tumour biology to be exploited therapeutically. However, high drug attrition rates remain a challenge in developing such improved treatments in ALL. High drug attrition has been attributed to preclinical models not being clinically translatable. Key barriers hindering bench to bedside translation when using complex in vivo models include species specificity, high financial costs, and lengthy experiments. Patient samples do not always engraft in mice. Samples that do engraft form successful xenograft models in 3-6 months at which point drug testing can be started which takes another 3-4 weeks.

A 3D BM-mesenchyme-leukaemia spheroid model will provide researchers with a biomimetic platform where clinical samples can be cultured within the context of their microenvironment, and patient specific drug testing performed within a week. Besides delivering translatable drug response data within clinically relevant timeframes the relative simplicity of ex vivo spheroid models mean that they are tractable, transferrable and sustainable. Consequently, such models have wide applications in translational research in academia and industry alike. These models can be set up in laboratories and SMEs locally, nationally and globally that do not contain infrastructure for animal procedures. Following further extensive validation, such spheroid biobanks also have the potential to be embedded within clinical trials and healthcare systems for the purposes of detecting responders as well as to aid risk stratification.

In this paper we show that our 2D model itself brought about significant animal replacement locally. Leukaemia research at Newcastle requires 3600 animal procedures every year. Local metrics confirmed that using the 2D pilot approach we replaced mice in 67 drug tests last year. Given minimum of 40 animals are needed per drug test we replaced 2680 animals (out of 3600) which constituted a minimum 73% local animal replacement. To improve biomimicry and consequently translatability and transferability of our model, we develop 3D BM-mesenchyme spheroids. We show that these 3D spheroids show superior ability in supporting culture of leukaemia patient samples. The spheroids also provide improved treatment protection when compared to 2D co-cultures and therefore are appropriate in modelling treatment resistance which remains a major clinical challenge in cancer management. This simple and tractable 3D prototype will form the first steppingstone in developing next generation 3D preclinical models with improved in vivo and patient drug response predictability. Such sustainable ex vivo models will ultimately replace existing moderate severity procedures in leukaemia research with models that successfully impact clinical outcome.

## ACKNOWLEDGEMENTS

This study was funded by NC3Rs grants to DP. We thank Professor Olaf Heidenreich and Dr. Kenneth Rankin, Newcastle University for provision of samples.

## FUNDING

National Centre for the Replacement Refinement and Reduction of Animals in Research (NC3Rs) - NC/P002412/1 [DP]

National Centre for the Replacement Refinement and Reduction of Animals in Research (NC3Rs) - NC/V001639/1 [DP]

## AUTHOR CONTRIBUTIONS

Conceptualization: DP

Methodology: DP, HB, AW

Investigation: AW, SH, JP, HB, DP

Visualization: AW, SH, DP, HB, JP

Resources: DP, HB

Funding acquisition: DP

Project administration: DP, HB

Supervision: DP, HB

Writing – original draft: DP, AW

Writing – review & editing: DP, AW, SH, JP, HB

